# Local context influences memory for emotional stimuli but not electrophysiological markers of emotion-dependent attention

**DOI:** 10.1101/184309

**Authors:** Gemma E. Barnacle, Dimitris Tsivilis, Alexandre Schaefer, Deborah Talmi

## Abstract

Emotional Enhancement of free recall can be context dependent. It is readily observed when emotional and neutral scenes are encoded and recalled together, in a ‘mixed’ list, but diminishes when these scenes are encoded separately, in ‘pure’ lists. We examined the hypothesis that this effect is due to differences of allocation of attention to neutral stimuli according to whether they are presented in mixed or pure lists, especially when encoding is intentional. Using picture stimuli that were controlled for semantic relatedness, our results contradicted this hypothesis. The amplitude of well-known electrophysiological markers of emotion-related attention - the Early Posterior Negativity (EPN), the Late Positive Potential (LPP), and the Slow Wave (SW) - was higher for emotional stimuli. Crucially, the emotional modulation of these ERPs was insensitive to list context, observed equally in pure and mixed lists. Although list context did not modulate neural markers of emotion-related attention, list context did modulate the effect of emotion on free recall. The apparent decoupling of the emotional effects on attention and memory challenges existing hypotheses accounting for the emotional enhancement of memory. We close by discussing whether findings are more compatible with an alternative hypothesis, where the magnitude of emotional memory enhancement is, at least in part, a consequence of retrieval dynamics.

## Introduction

Scenes that depict accidents, violence, and war trigger intense negative feelings and capture our attention involuntarily. The fact that people are able to retrieve the gist of such scenes and describe them later, and that this ability is superior for emotional than for neutral scenes (Dolcos, Denkova, & Dolcos, 2012; Talmi, 2013), is therefore not particularly astonishing. Indeed, emotional enhancement of memory is thought to be evolutionarily adaptive, in that we remember best those events that are important to us, namely, those that triggered us to respond emotionally at encoding. Because the effect of emotion on memory is thought to be adaptive, it is intriguing to observe that it can be context-dependent. Behavioural work has established that emotional memory enhancement in free recall tests of early long-term memory is readily observed when emotional and neutral stimuli are encoded and recalled together in ‘mixed’ lists but is weaker when these scenes are encoded separately, in ‘pure’ lists, and can sometimes disappears completely when pure lists are additionally controlled for confounding factors such as differential organisation and attention (Barnacle et al., 2016; Hadley & MacKay, 2006; Sommer et al., 2008; Talmi et al., 2012; Talmi et al., 2007; Talmi & McGarry, 2012). The context dependence of the effect of emotion on memory is surprising because a pure list of emotional scenes is effectively an operationalisation of a real-life event that consists of a number of emotional aspects; for example, witnessing a traffic accident where one might observe injured persons, damage to property, medical personnel and so on. The evolutionary logic would predict that aspects of an emotional event would be remembered better than aspects of neutral events.

Behavioural experiments that employed the divided-attention paradigm have established that emotional scenes presented in mixed lists capture attention preferentially (Kensinger & Corkin, 2004; Kern et al., 2005; Talmi, 2013). It has been argued, therefore, that enhanced attention to emotional stimuli at the expense of attention to temporally or spatially adjacent neutral stimuli could explain the emotional enhancement of memory in tests of early long-term memory (Hamann, 2001; Mather & Knight, 2009; Mather & Sutherland, 2011). For the temporal effects of emotional stimuli, a series of studies concluded that direct resource competition between emotional and neutral words is only observed when the inter-trial intervals (ITIs) are short (up to two seconds, Schmidt & Schmidt, 2016). Still, when the stimuli are more complex, such as the emotional scenes that are widely employed in the emotional memory literature, it is possible that such competition may still be present even with longer ITIs.

If the attentional advantage of emotional stimuli is reduced in pure lists, perhaps because emotional stimuli are expected in that list’s context (Barrett & Bar, 2009), their memory advantage may be reduced as a direct consequence. The behavioural evidence for this hypothesis is inconclusive. On the one hand there is evidence that emotional stimuli produce states of ‘vigilance’, detected through their influence on the perception of neutral stimuli (Golomb, Turk-Browne, & Chun, 2010). For example, emotional words influence the reading time and font-color naming time of neutral words presented in the same block (Algom et al., 2004; McKenna & Sharma, 2004; Schmidt & Saari, 2007). These findings show that emotional stimuli influence participants’ expectations about the type of stimulus they may encounter next, and could lead to reduced attention to highly expected emotional stimuli. On the other hand, there is also evidence that individual emotional stimuli in pure lists still attract extra processing resources compared to neutral stimuli in pure lists. For example, font-colour naming of a block of taboo words takes longer than font-colour naming of a block of neutral words. Similarly, performance on a secondary task was impaired equally when participants viewed emotional (compared to neutral) words or scenes both when those scenes were presented in mixed lists and when they were presented in pure lists (Schmidt & Saari, 2007; Talmi & McGarry, 2012). In evaluating the behavioural evidence, it is important to acknowledge that behavioural assays of attention may not be sufficiently sensitive to the dynamics of encoding, because they only collect discrete responses every few seconds. To overcome this limitation here we used EEG with the aim to illuminate how attention to emotional and neutral information is modulated by list composition.

Electrophysiological research has established which ERPs are modulated by the emotionality of stimuli. Three ERPs, including the Early Posterior Negativity (EPN), the LPP, and the Slow Wave (SW), are known to be sensitive to visual attention and are robustly modulated by emotion (Schupp et al., 2006). These three ERPs are thought to index different processes of emotion-guided selective attention (Foti et al., 2009; Schupp et al., 2006). The LPP, in particular, has been shown to index emotionally-biased attention even when potential confounding visual differences between emotional and neutral stimuli were eliminated, by comparing neutral objects that were previously paired with emotional or neutral contexts (Ventura-Bort, Low, et al., 2016). Our research question hinges on whether these ERPs are modulated by the local list context. Some results suggest that the LPP is sensitive to incongruity (Herring et al., 2011), which supports the suggestion that this component will be affected by list context. Yet previous results that examined this question directly using series of emotional and neutral scenes are somewhat inconclusive. Schupp et al. (2012) found that the valence of the preceding sequence of pictures did not attenuate the emotional modulation of the EPN and the LPP to the final picture, supporting the contextual independence of these ERPs. Similarly, in two studies, Codispoti and colleagues found that the emotional modulation of the LPP remained intact even when the same emotional and neutral pictures were presented up to 60 times (Codispoti et al., 2006, 2007). Pastor et al. 2008) did find evidence of contextual dependence, in that the emotional modulation of the SW was stronger in pure lists compared to mixed lists over frontocentral and occipital electrodes, and the LPP associated with neutral pictures over occipital electrodes was also affected by list context. But the dependence Pastor et al. (2008) observed did not comply with the logic of the hypothesis we propose to test here, that the emotional modulation of the relevant ERPs would be reduced in pure lists compared to mixed lists. Crucially, these studies used orienting tasks, such as passive viewing or emotionality ratings, which do not give participants reasons to pay special attention to neutral stimuli. Perhaps when participants know that their memory would be tested they attempt to pay attention to all stimuli, but fail when presented with neutral stimuli in mixed lists because of competition for resources. If correct, we should find that list context modulates the emotional modulation of the EPN, LPP and SW in intentional encoding conditions.

Dolcos and Cabeza (2002) were the first to investigate the electrophysiological correlates of the emotional enhancement of memory for scenes. They observed subsequent memory effects for emotional stimuli both early (400-600ms) and late (600-800ms), but only a late effect for neutral stimuli. The exact timing of the emotion-by-memory interaction varied in subsequent studies (Righi et al., 2012; Weymar, Löw, Melzig, & Hamm, 2009), but Dolcos and Cabeza’s conclusion that emotional scenes have privileged access to mnemonic resources at encoding was supported, and was one of the motivations for our current hypothesis that attention allocation must be a key factor for the context-dependence of emotional enhancement of memory. Only one previous electrophysiological study has manipulated both emotion and context in the context of a memory task (Watts et al., 2014). Behaviourally, Watts and colleagues observed a stronger emotional enhancement of memory for mixed compared to pure lists, although the effect was statistically significant for both list types. Their ERP data showed that the subsequent memory effect (or the “Dm” effect, Paller & Wagner, 2002) for neutral pictures in posterior sites was reduced in mixed lists compared to pure lists in an early (200 to 400ms after picture onset) and a late (800-1500ms) time windows. The neural and the behavioural findings thus converged, and interpreted as suggesting a less efficient encoding of neutral pictures in mixed lists, in accord with our current research hypothesis. Another interpretation of these results is that encoding activity was not as good a predictor for neutral stimuli in mixed lists, compared to other conditions. We return to this alternative in the discussion section. Because Watts et al. (2014) focused on ERP correlates of subsequent memory (the Dm effect), they could not specifically examine the ERP correlates of emotion-related selective attention. Indeed, it is unclear if the LPP and EPN can be readily operationalized from Dm-related ERP activity. In addition, similarly to most studies with emotional pictures (including Dolcos & Cabeza, 2002; Pastor et al., 2008; Schupp et al., 2012), Watts and colleagues did not control their emotional and neutral stimuli for differences in semantic relatedness. Negative emotional stimuli are related thematically (e.g. a crime scene is related to a scene depicting a woman crying), and are therefore typically more cohesive than unselected neutral stimuli (Talmi & Moscovitch, 2004), a factor which may influence memory performance especially when the theoretical mechanism at play centres on what participants expect to see in pure or mixed list contexts. For example, we know that the violation of expectation can modulate free recall (Hirshman, 1988).

In summary, the list context effect on the emotional enhancement of memory is a robust behavioural effect, but its underlying mechanism is still not known. Previous work suggests that this effect is caused by an enhancement of selective attention towards emotional items in mixed but not pure lists, or perhaps diminished attention to neutral stimuli in mixed compared to pure lists when encoding is intentional. The aim of this study was to test this account using well-known electrophysiological markers of emotion-related attention. Here we examine the EPN, LPP and SW when participants encode pure and mixed lists, controlled for semantic relatedness. The emotional modulation of the EPN, which is thought to reflect early selection of stimulus processing, is unlikely to be affected by list context. We hypothesised, based on the previous work, that the emotional modulation of the LPP and the SW, which are thought to reflect the enhanced visual processing and sustained attention to emotional stimuli, would be heightened in mixed compared to pure lists, and perhaps even absent in pure lists, in close parallel with the free recall findings.

## Method

### Participants

Twenty-five healthy adults, age 18-35, with no current or past history of neurological or psychiatric illness were recruited through advertisements and the University of Manchester student credit participation system. Two participants were excluded because of technical failures during the recording. As described below, if any of the channels had more than 20% bad trials that channel was excluded from analysis; participants who had more than one bad channel of those selected for analysis were excluded altogether. This approach lead us to exclude one additional participant, leaving a final sample of N=22. Three additional participants had more than one bad channel of those selected for the analysis of EPN, and they were excluded only from that analysis (leaving a subsample of N=19 for the EPN analysis). Participants provided informed consent and were reimbursed for their time and expenses by course participation credits or £15. Ethical approval was obtained from the University of Manchester Research Ethics Committee.

### Materials and Equipment

Experimental stimuli consisted of 238 colour images (size: 280x210 pixels), half of which conveyed negative valence and were arousing (hereafter referred to as “emotional”), and half of which were neutral in valence and not arousing (hereafter referred to as “neutral”). Participants judged emotional stimuli to be semantically related to each other (e.g. (Talmi & Moscovitch, 2004)). In order to control for this factor within the neutral set the theme of ‘domesticity’ was chosen such that all neutral pictures depicted domestic scenes. All experimental stimuli contained at least one human being. Of the total pictures, 14 were practice pictures (displayed only in the practice block), and 32 were buffer pictures (16 neutral and 16 emotional); both were excluded from behavioural and EEG analysis. The pictures were taken from the internet and IAPS pictures (Lang, Bradley, & Cuthbert, 2008) and were rated in a separate study where participants rated the pictures for arousal and valence in one session, and for semantic relatedness in another session; session order was counterbalanced. Arousal and valence were rated using the self-assessment manikin arousal and valence scales (Bradley & Lang, 1994). Semantic relatedness was rated on a 1-7 scale, where ‘7’ indicated that the target picture was closely related to a ‘standard’ set of 9 pictures of the same valence. The standard sets (one emotional, one neutral) broadly represented the content of the entire stimulus pool of the same valence.

The emotional and neutral stimuli selected for use in the current experiment were significantly different on measures of arousal, *t*(31) = 13.80, *p* < .001, *ŋ*^2^ = 0.75; valence, *t*(31) = 15.46, *p* < .001, *ŋ^2^* = 0.79; and were equated for measures of semantic relatedness, *t*(27) = 1.32, *p* = .20, *ŋ*^2^ = 0.03; see Table 1.

**Table 1.**

|  | Neutral |  | Emotional |  |
| --- | --- | --- | --- | --- |
|  | *M* | *SD* | *M* | *SD* |
| Arousal* | 2.34 | 1.36 | 5.65 | 1.41 |
| Valence* | 5.34 | 0.49 | 2.81 | 0.66 |
| Semantic Relatedness | 5.22 | 1.31 | 4.81 | 1.24 |

*Note.* Mean *(M)* and standard deviation *(SD)* statistics from ratings of all experimental pictures. Arousal scale 1-9: (1=low arousal, 9=high arousal); Valence scale: 1-9 (1=negative, 9=positive); Semantic relatedness scale: 1-7 (1=low relatedness, 7=high relatedness). * indicates measures were significantly different from between valence categories.

Stimuli were randomly allocated to 16 experimental lists: 8 mixed lists and 8 pure lists (4 pure lists of each valence). Mixed lists contained two buffer stimuli (one of each valence, randomised in order of presentation) presented at the beginning of each list, and excluded from subsequent analyses to reduce the impact of primacy effects; followed by 12 stimuli - 6 from the emotional and 6 from the neutral sets, in a randomised order. Pure lists contained two same-valence buffer stimuli followed by 12 same-valence stimuli (either all neutral or all emotional). The allocation of stimulus to list type, the order of lists presented, and the order of stimuli within lists were randomised.

Stimuli were displayed on a 15” by 12” screen, which was positioned approximately 95cm from the participant. Stimulus presentation and programming was realised using Cogent 2000 (Wellcome Department of Imaging Neuroscience, UCL, UK; http://www.vislab.ucl.ac.uk/cogent_2000.php).

### Procedure

Our procedure resembled that used by Talmi & McGarry (2012) and Dolcos & Cabeza (2002). Each participant undertook one practice block and sixteen experimental blocks. Each block included three tasks: list encoding, distractor, and free recall. Instructions were presented to the participant for each task on screen, and read aloud by the experimenter at the beginning of the experiment. Participants performed the encoding and distractor tasks alone in the room. Immediately after this, the experimenter re-entered the room in order to record the participant’s free recall responses. In deviation from (Talmi & McGarry, 2012), the experimenter wrote down the responses as they were spoken by the participant to reduce movement of the EEG head cap that can influence the EEG measurements in the next block. EEG was recorded throughout list encoding, but not during distractor and free recall. The experimenter monitored eye movement artefacts in real-time by observing the continuous EEG data during the recording. Feedback was given to participants if they were not conforming to the instructions to remain still, fixate on the cross and withhold blinks whilst the stimuli were displayed. These instructions were tolerated well by all participants after the practice block.

#### List Encoding

In each block participants passively encoded one list of pictures under intentional encoding instructions. A fixation cross was presented 500ms before each picture was displayed and remained on the screen overlaid on the image, which helped to prevent saccadic eye movements (participants were instructed to focus on the fixation and suppress eye movements). Each picture was presented for 2000ms with a jittered inter stimulus interval of 4000ms +/-500ms. This long ITI was chosen, following (Talmi & McGarry, 2012), to eliminate carry-over effects (Schmidt & Schmidt, 2016).

#### Distractor Task

After viewing the pictures participants engaged in an arithmetic task, which aimed to eliminate the contribution of working memory to the recall output. Two simple sums were presented, one each on the right and left hand side of the screen. Participants were asked to compute the sums mentally and identify the highest value sum using two keys relating to the right or the left of the screen (‘2’ for the left and ‘3’ for the right, using the number keypad on the keyboard). A keyboard placed in front of the participant within comfortable reach allowed the participant to make their selections when prompted. The distractor task lasted for sixty seconds, after which the words ‘free recall’ were presented on screen.

#### Free Recall Task

The experimenter re-entered the EEG chamber and asked participants to recall as many pictures from the previous list as they could remember, in any order and in as much detail as possible. Participants were asked to be specific in their descriptions of stimuli such that descriptions of two similar pictures should be distinguishable from their responses. Participants were given 3 minutes for this task.

### EEG Recording and data reduction

BioSemi Active Two measurement system (BioSemi, Amsterdam, www.biosemi.com) was used to measure EEG activity from the scalp using 64 electrodes and conforming to the 10-20 system embedded in an elasticated cap (Chatrian, Lettich, & Nelson, 1985). This system allows for high-input impedance, and thus classical impedance thresholds do not apply, and the classical measurement of impedance is not feasible (E. S. Kappenman & Luck, 2010); impedance information was not recorded. VEOG electrodes were used for detecting eye artefacts. The EEG signal was recorded using Actiview software, which applies a 0.16Hz on-line highpass filter and a 100Hz on-line lowpass filter. Data was pre-processed using SPM8 (www.fil.ion.ucl.ac.uk/spm/) (Litvak et al., 2011). The data was re-referenced offline to the combined mastoids reference, thought to optimise LPP effects (Hajcak et al., 2012), filtered between 0.1 and 25 Hz, down-sampled to 125 Hz, and epoched between -200 and +5500ms time-locked to stimulus onset. Individual participants’ eyeblinks were identified in their continuous data, and an epoch identified from -500ms to +500ms relative to the peak of the blink. An average of the eye-blink topography per participant was then created using the singular value decomposition (SVD) method, and this data was then removed from the epoched EEG using the signal source projection method (SSP, Nolte & Hämäläinen, 2001). An artefact rejection threshold of 250mV was applied before this procedure and a second artefact rejection threshold of 120mV applied after this procedure. Remaining trials were then averaged using the robust averaging algorithm (Litvak et al., 2010), a method that down-weights outliers. Averaged data was filtered again with a low-pass filter of 25Hz to remove any noise introduced from the process of robust averaging procedure (following standard practice, see user manual) and baseline-corrected. The mean amplitudes of the ERPs were extracted for each participant to represent the EPN, LPP and slow wave. The time window for the EPN (150-350ms), the LPP (400-700ms) and the slow wave (1-5.5s) were based on Schupp et al. (2006, 2012). Previous work showed that the modulation of the SW by emotion lasts at least up to one second after picture offset (Hajcak & Olvet, 2008). We therefore parcelled out the SW to 4 time periods: the last second of picture presentation, (1-2s from picture onset), early ITI (2-3s from picture onset, namely 0-1s from picture offset), middle ITI (3-4s from onset; 1-2s from offset), and late ITI (4-5.5s from onset; 2-3.5s from offset). Again following Schupp et al. (2006, 2012), the EPN was averaged across electrodes Oz, POz, O1, O2, PO3, PO4, PO7, PO8 and the LPP and slow wave were averaged across centro-parietal electrodes Cz, CPz, Pz, C1, C2, P1, P2, CP1, CP2. An additional analysis of trials that were later remembered was conducted on a subsample of participants who contributed more than 12 such trials in every condition (N=13 in the LPP/SW analysis and N=12 in the EPN analysis).

For completion, we have extracted peak amplitudes for the N1, N2, and P3, separately for each condition (Olofsson, Nordin, Sequeira, & Polich, 2008). We have extracted the negative peaks for N1 and N2 from the aggregate EPN electrodes at 150-200ms and 200- 350ms, respectively, for the N=19 participants for whom we reported results of the EPN component, and positive peak for P3 from the average of Pz, P1 and P2 electrodes for the complete N=22 sample for whom we reported results of the LPP component.

All statistical analysis was conducted in SPSS version 22 (IBM analytics). Greenhouse-Geisser correction was applied when necessary. A significance threshold of p<.05 was used throughout; non-significant results with p<.10 are also reported in full. Bonferroni-corrected t-tests were used to unpack significant interactions whilst controlling for multiple comparisons.

## Results

### Behavioural Results

Free recall responses were scored following previous work (Bradley, Greenwald, Petry, & Lang, 1992; Talmi &McGarry, 2012). The experimenter matched the participant’s descriptions of the pictures seen in each block to the experimental stimuli seen in that block. Recall responses were coded by a second independent coder, and agreement amongst coders was high (97%). Disagreements were resolved through discussion. Proportion scores - the number of correctly recalled items in a given condition divided by the total number of items of that kind seen in that condition - were entered into a 2 (list composition: pure/mixed) x 2 (emotion: negative/neutral) repeated measures ANOVA. This analysis identified a significant main effect of emotion, F(1, 21) = 28.50, *p <* .001, ŋ_p_^2^ = .58). The effect of list context did not reach significance, F(1,21)=4.06, p=.057, ŋ_p_^2^=.16. As expected, there was a significant emotion by list context interaction F(1, 21) = 33.12, *p* < .001, ŋ_p_^2^ = .61, and post-hoc tests revealed that emotional enhancement of memory was significant and large in mixed lists, *t*(22) = 6.90, *p* < .001, ŋ^2^= 0.35), but not significant and small in pure lists *t*(22) = 1.72, *p =* .10, ŋ^2^= 0.03 (see Figure 1). Significantly fewer neutral stimuli were remembered in mixed compared to pure lists *t*(22) = 4.96, *p <* .001, ŋ^2^= 0.22), while more emotional stimuli were recalled in mixed compared to pure lists *t*(22) = 2.93, *p* = .01, ŋ^2^= 0.09).

**Figure 1.**
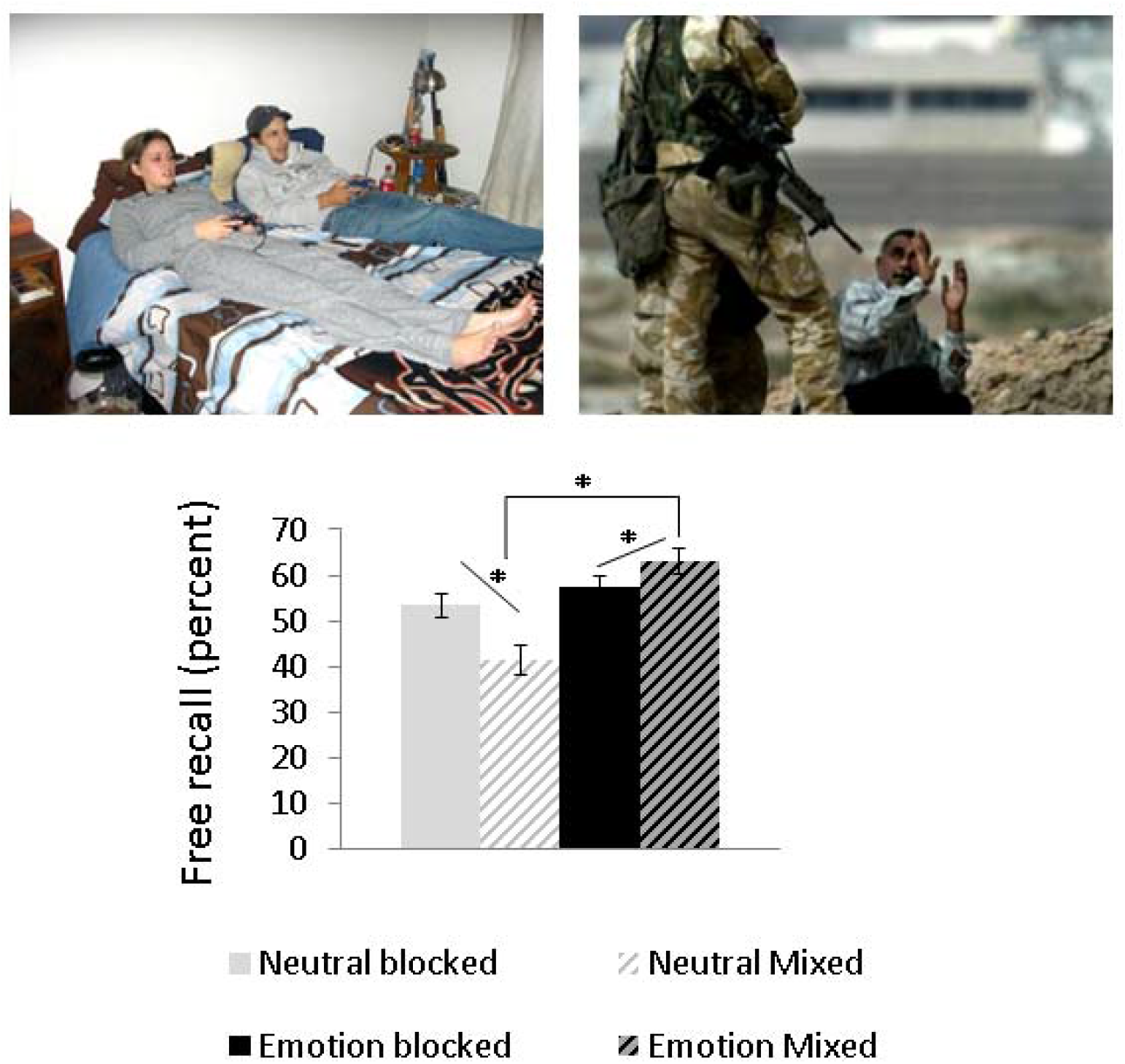
Proportion recall. Average proportion recalled items in pure and mixed conditions for emotional and neutral stimuli. The asterisk designates significant effects. Error bars indicate standard error.

### EEG Results

Participants contributed an average of 39.78 artefact-free trials (SD=4.07) for every relevant trial type. In each condition we analysed had 37 artefact-free trials on average, with each participant contributing at least 21 artefact-free trials in each condition. The subsample of N=19 participants who were included in the analysis of EPN provided an average of 40.44 artefact-free trials (SD=3.98). Adding the factor of memory status (recalled vs. forgotten) reduced the number of trials per cell substantially, so we could not justify an analysis of subsequent memory effects. We therefore limited ourselves to describing emotional modulation of attention regardless of subsequent memory. Extracted data corresponding to the EPN, LPP and SW were entered to separate 2 (list context: pure/mixed) x 2 (emotion: negative/neutral) repeated measures ANOVAs.

#### EPN

Extracted data from occipitoparietal electrodes were entered into a repeated-measures ANOVA with the same factors as above (Figure 2). In accordance with previous literature on this component, the amplitude of the EPN was significantly higher for emotional than neutral scenes with a medium effect size, F(1,18) = 6.75, *p =* .18, ŋ_p_^2^ = .27. Neither the effect of list context nor its interaction with emotion was significant. The subsample analysis of hits qualitatively replicated these results, but here the effect of emotion was only present at a trend level, although it remained an effect of medium size, F(1,11) = 3.33, *p* = 0.09, ŋ_p_^2^ = .23. Other effects were not significant.

**Figure 2.**
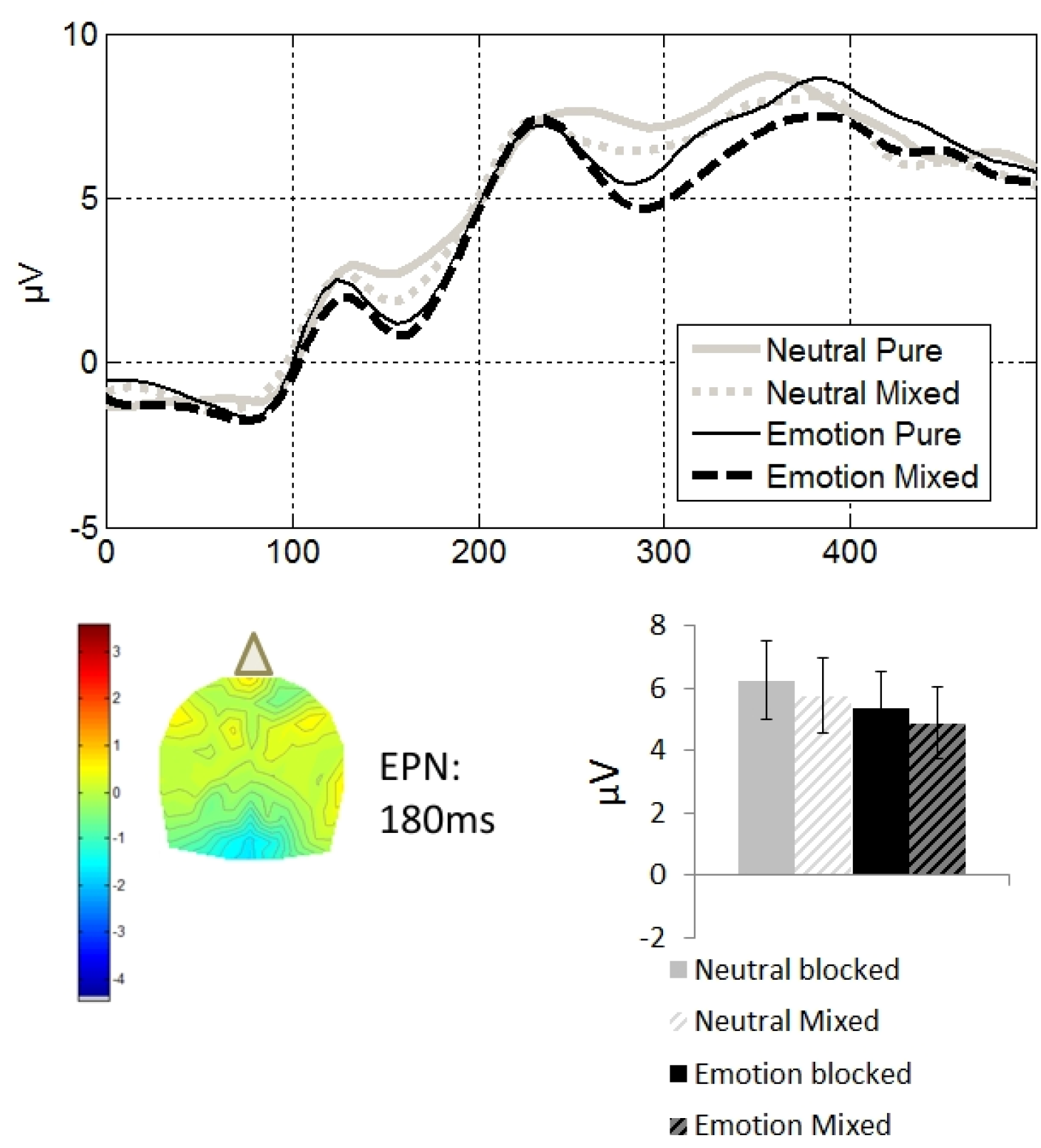
Occipitoparietal effects as a function of list context (solid: pure lists; hashed: mixed lists) and emotion (black: emotional, grey: neutral). Data were time-locked to picture presentation and extracted from occipitoparietal electrodes Oz, POz, O1, O2, PO3, PO4, PO7, and PO8. Top: ERPs traces depicting the EPN, N1 and N2. Bottom left: 2D topographical maps of the latency in which the difference between the signal in the emotional and the neutral conditions was maximal within the time window of the EPN, 180ms from picture presentation, collapsing across pure and mixed lists. The top and bottom of the topographies correspond to the front and the back of the head. Bottom right: average amplitudes across the time windows corresponding to the EPN, 150-3 50ms from picture presentation. Error bars indicate standard error.

#### LPP

Extracted data from centroparietal electrodes were entered into a repeated-measures ANOVA with the same factors as above (Figure 2). Emotion increased the amplitude of the LPP, F(1,21) = 11.58, *p* =.003, *ŋ_p_^2^* = .35. Neither the effect of list context, F(1,21) = 3.49, *p* =.08, *ŋ_p_^2^* = .14 nor its interaction with emotion reached significance. The subsample analysis of hits replicated these results with a significant and large effect of emotion, F(1,12) = 5.51, p = 037, ŋ_p_^2^ = .31. Other effects were not significant.

#### Slow Wave

The mixed emotion condition produced the highest amplitude, while amplitude in the mixed neutral condition was suppressed relative to all other conditions, an effect that was more pronounced earlier on in the SW epoch. Extracted data from centroparietal electrodes were entered into a 4x2x2 repeated-measures ANOVA with the factors time bin (picture present, early-, middle- and late-ITI), list context (mixed/pure) and emotion (emotional/neutral). The results are depicted in Figure 2. Unsurprisingly, the factor of time bin was significant, F(3,63) = 8.40, *p* =.001, as the component decreases with time. The trend for a main effect of emotion did not reach significance, F(1,21) = 3.81, *p* = .06, ŋ_p_^2^ = .15 but interacted significantly with time bin, F(3,63) = 4.18, *p* = .009, *Up* = .16, because it was stronger earlier in the epoch. No other effects were significant. To query the duration of the effect of emotion we conducted separate repeated-measures ANOVAs for each of the four time windows, with the factors of emotion and context. As expected, during the last second of picture presentation the main effect of emotion was significant, F(1,21) = 11.46, *p* =.003, ŋ_p_^2^ =.35; the other effects were not significant. The effect of emotion was no longer significant in the first second after picture offset (early ITI), F(1,21) = 3.98, *p* =.06, ŋ_p_^2^=. 16, nor at any other subsequent time bin. The subsample analysis of hits replicated the obvious effects of time, F(3,36) =10.63, *p* =.001, ŋ_p_^2^=.47, and here the main effect of emotion was significant, F(1,12) = 6.31, *p* =.027, ŋ_p_^2^ =.34, but these two effects did not interact significantly and the effect size of the interaction was small. This result implies that when events that are eventually remembered are considered, separately from those that will subsequently be forgotten, emotion appears to influence scalp markers of attention all throughout the ITI, but given the small sample size we should be cautious in interpreting the null interaction effect.

#### Correlations with EPN, LPP and Slow Wave

We examined correlations, in mixed and pure lists separately, between memory and ERPs - namely EPN, LPP and (average) SW. The three correlations within each list context were corrected for multiple comparisons (corrected *p* = .016). None of the correlations were significant. We note that cross-participants correlations in a small sample such as the one used here is likely underpowered.

#### Other components

The pattern that emerged when we examined additional components resembled the pattern that is already reported above (Figure 3). Peak amplitudes for the N1, N2 and P3 were analysed with a 2 (list context: pure/mixed) x 2 (emotion: negative/neutral) repeated measures ANOVA. Each of these analyses revealed a main effect of emotion (N1: F(1,18)=12.96, p=0.002, ¾^2^=.42; N2: F(1,18)=6.32, p=.022, ¾^2^=.26; P3: F(1,21)=6.75, p=.017, ¾^2^=.24). For the P3, there was a trend towards a more positive amplitude in pure, compared to mixed lists (F1,21)=3.86,p=0.06). No other effects were significant.

**Figure 3.**
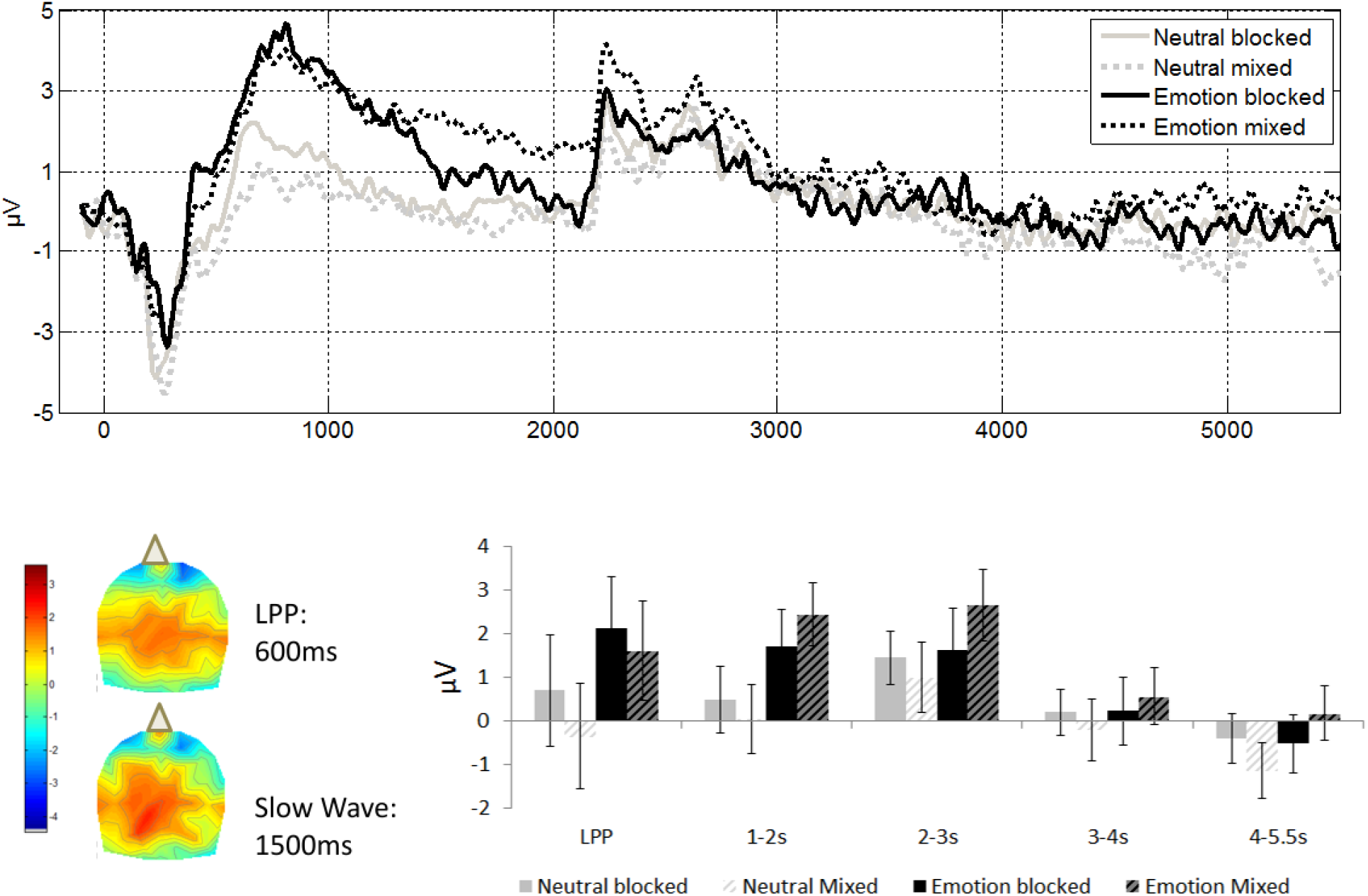
Centroparietal effects as a function of list context (solid: pure lists; hashed: mixed lists) and emotion (black: emotional, grey: neutral). Data were time-locked to picture presentation and extracted from centroparietal electrodes Cz, CPz, Pz, C1, C2, P1, P2, CP1, and CP2. Top: ERPs traces depicting the LPP, Slow Wave and the P3. Middle panels: 2D topographical maps of the latency in which the difference between the signal in the emotional and the neutral conditions was maximal within the time window of the LPP, 600ms from picture presentation, and the Slow Wave, 1500ms from picture presentation, collapsing across pure and mixed lists. The top and bottom of the topographies correspond to the front and the back of the head. Bottom right: average amplitudes across the time windows corresponding to the LPP (400-700ms from picture presentation) and slow Wave (1000-5500ms, broken down to 4 time bins as indicated on the x-axis). Error bars indicate standard error.

## Discussion

This study tested the hypothesis that the effect of list context on behavioural emotional memory performance is caused by an enhancement of selective attention towards emotional stimuli only in mixed lists but not in pure lists. We tested this hypothesis by examining the most widely known ERP correlates of motivated attention. The behavioural results replicated our previous work, demonstrating an emotional enhancement of memory in mixed but not pure lists. As we have observed previously, memory for neutral stimuli in mixed lists was decreased (a large effect size), and contributed to the interaction between list context and emotion, together with the increase in memory for emotional stimuli in mixed lists (a small effect size), a more subtle effect which is not always observed. In contradiction with our hypothesis, whereas list context modulated the effects of emotion on behavioural memory performance, it did not modulate the effects of emotion on neural indicators of emotion-related attention at encoding. Replicating previous findings, emotion modulated the EPN, LPP and the SW, which are established correlates of emotion-related attention, as well as the N2, N2 and P3. Crucially, the emotional modulation of these ERPs was context-independent, and statistically they were equally strong in pure and mixed lists. The same findings were obtained in an analysis that focused on subsequently remembered items, although we acknowledge that a more powered analysis of hits may expose context effects. These results refute both the suggestion that attention is no longer preferentially allocated to emotional stimuli in pure lists, and the suggestion that attention to neutral stimuli in mixed lists is severely depleted compared to attention to the same pictures in pure lists. List context here effectively created a functional dissociation between attention (indexed neurally) and memory (indexed behaviourally). These available results are intriguing because they impose constraints on models suggesting that additional attention to emotional items at encoding is the main determinant of emotional memory enhancement. We turn to these theoretical implications after we discuss the electrophysiological results in more detail.

The finding that emotion modulated the EPN, LPP and SW is important because this is the first study to examine the effect of emotion on these components during an intentional encoding task, in which participants have a reason to pay attention to neutral stimuli, and the first time the comparison neutral scenes were controlled for semantic cohesiveness. The emotion modulation of the LPP and SW is typically considered to be a result of bottom-up modulation of visual attention; but when participants intentionally encode neutral items for a subsequent test they could well recruit visual attentional resources through top-down means. Indeed, in our recent fMRI study the successful encoding of neutral scenes in mixed lists relied on top-down attentional resources to a greater extent compared to the successful encoding of neutral scenes in pure lists (Barnacle et al., 2016). It was therefore entirely possible that the emotional modulation of these ERPs would be attenuated in our task, where participants allocated their full attention to intentional encoding (Holmes et al., 2014). This is especially true for the pure-list condition, where memory for emotional and neutral stimuli was equivalent, and where there was no competition from neighbouring emotional items. Our findings that emotion modulated these ERPs even in an intentional encoding task and regardless of local list context therefore support conclusions that the effect of emotion is obligatory and independent of task demands (Codispoti et al., 2006; Schupp et al., 2012).

Our electrophysiological results may be seen as contradicting those of Pastor et al. (2008) who observed that the LPP and the SW were modulated by list context. A potentially important difference between Pastor et al.’s study and the current study is that the current study controlled for the semantic relatedness of emotional and neutral stimuli, whereas previous studies did not. Stimuli drawn from a semantically cohesive category would be less incongruent with other stimuli, a factor known to influence the amplitude of the LPP (Herring et al., 2011). Furthermore, semantic cohesiveness is thought to be a driver for the mobilization of attentional processes. Specifically, when stimuli are presented in sequential lists their semantic relatedness with other items in the list may engage attentional resources that are be used for encoding strategies based on inter-item relatedness (Watts et al., 2014). In support of this idea, Dillon et al. (2008) found that a frontal slow positive wave (up to 700ms) for emotional stimuli could be explained by semantic cohesiveness. Also, others (Otten, Sveen, & Quayle, 2007; Paller & Wagner, 2002) have suggested that a positivity between 400 and 1000ms post-stimulus onset for subsequently remembered items reflected an enhanced processing of semantic features of items at encoding. As suggested by Watts et al. (2014), emotional stimuli can be prioritised in mixed encoding lists because they provide opportunities for encoding strategies based on semantic relatedness that neutral stimuli do not. These potential prioritisation strategies would not be feasible when semantic interrelatedness is equated between different stimuli types included in the encoding lists, as in the current study. Therefore, it may be possible that when semantic cohesiveness is unconstrained, the increased semantic cohesiveness of emotional stimuli would modulate attentional processes, resulting in context effects; indeed, the lack of control over this factor in previous research could explain the differences between Pastor et al.'s 2008) and Schupp et al. (2012).

Our results also touch on the question of how the emotional response evolves during the entire experimental session. First, the results are relevant to the question of whether the emotional response habituates during the presentation of pure emotional lists. This is important because such habituation could explain why emotion does not enhance memory in the pure list condition. Previous results showed that some habituation does occur in experiments with emotional stimuli, but it is far from complete. For example, Bradley, Lang and Cuthbert (1993) presented the same 6 pleasant, unpleasant and neutral pictures repeatedly and observed habituation in heart rate, skin conductance responses, and electromyography measures over time. However, if we examine their first block of trials, emotional and neutral pictures were differentiated in all of these measurements even though participants saw the same 6 pictures 4 times each (a total of 24 pictures). Later in that experiment, after 12 presentations of each picture, Bradley and colleagues still observed an emotional modulation of another physiological index of arousal, the startle response. Similarly, Pastor et al. (2008) also observed differential effects of habituation in two physiological markers of emotional arousal: increased habituation of the emotion modulation of the heart rate but decreased habituation of the emotion modulation of the skin conductance response in pure lists compared to mixed lists. Taken together, previous work suggests that although some indices of the emotional responses do habituate, others remain observable even after many presentations of emotional scenes. Notably, we used shorter lists than those that Bradley et al. have used, and - unlike Bradley et al. - used trial-unique pictures, which were all quite different from each other (e.g. a dead dog; a gunpoint), so we did not expect pronounced habituation to take place in this study (Rankin et al., 2009). In agreement, the emotional effect on ERP markers of attention in pure lists was not weaker than the effect in mixed lists.

A second aspect of the evolution of the emotional response in our study has to do with the predications of Arousal-Biased Competition theory (ABC theory, Mather & Sutherland, 2011) for our task. ABC theory proposes that arousing stimuli are assigned higher priority than neutral stimuli, and thereby garner additional processing resources. If the presence of emotional stimuli in mixed lists renders the entire mixed list context more arousing than the context of encoding of a pure list of neutral stimuli, the prioritisation of emotional stimuli would be amplified in this condition at the expense of neutral stimuli (Mather & Sutherland, 2011), because emotional stimuli would capture additional attention resources. In our task, such mechanisms would have been evident in higher ERP amplitudes in the emotion mixed compared to the emotion pure condition, and lower ERP amplitudes in the neutral mixed compared to the neutral pure condition. By contrast, none of the ERPs we examined, either emotional or neutral, differed in amplitude as a function of list context. In order to reconcile these findings with ABC theory we may hypothesize that the entire experimental session may have been more arousing as a result of including some emotional pictures, something that can be tested in future research with a between-subject design. More generally, our study shows the benefit that electrophysiological work can have for constraining the predictions of ABC theory when applied to novel tasks (Barnacle & Talmi, in press).

### Theoretical implications

Before we turn to the theoretical implications of our findings we note that while the functional dissociation between the effect of emotion on attention and memory contradicted our initial hypothesis, the dissociation is not entirely unexpected. In fact, these results add to mounting evidence that the emotional enhancement of memory may not depend only on encoding dynamics. For example, in our previous work, we showed that arousal had direct effects on early long-term free recall tests of memory even when attentional effects were covaried out statistically (Pottage & Schaefer, 2012; Talmi, Ziegler, et al., 2012; Talmi & McGarry, 2012) or through experimental manipulation using divided-attention (Talmi & McGarry, 2012). In our fMRI study, which used an identical paradigm to the one reported here, we have also not observed any evidence for reduced attention to neutral stimuli in mixed lists (Barnacle et al., 2016). How should we understand the decoupling of attention and memory, which are typically closely linked? Why do participants who pay more attention to emotional pictures in pure lists not remember them better than neutral pictures?

To answer these questions we developed a temporal context model of emotional memory enhancement, called the Emotional Context Maintenance Model (eCMR, Talmi, Lohnas & Daw, in preparation). The model relies on the fundamental understanding that when the recall context matches the encoding context, the match renders stimuli more retrievable; this has been demonstrated for temporal, semantic, and task contexts (Polyn et al., 2009), while context mismatches have profound effects on recall (Jonker, Seli, & MacLeod, 2013). Because emotional stimuli are preferentially attended they are better equipped to win the competition for retrieval at the time of test. This makes them more likely to be recalled earlier than less-well attended neutral items. In support of this proposal, when participants recall mixed lists they output emotional stimuli sooner than neutral stimuli (Talmi et al., 2007). According to temporal context models, whenever a stimulus is recalled, it retrieves its context too. Our model refers to an emotional context, building on the understanding that emotional scenes trigger systemic arousal that can last for many minutes (e.g. Henckens, van Wingen, Joels, & Fernandez, 2012). We propose, therefore, that an emotional stimulus retrieves its emotional context together with its temporal context, thereby helping the recall of further emotional items that share the same emotional context, while hindering recall of stimuli with a different (neutral) context (Polyn et al., 2009). This interpretation is directly supported by evidence that objects that have been paired just once with an emotional context are associated with enhanced LPP magnitude and enhanced old-new effects during a subsequent recognition test (Ventura-Bort, Low, et al., 2016; Ventura-Bort, Löw, et al., 2016). Further, this interpretation is also supported by findings that participants who recall mixed lists of emotional and neutral stimuli tend to recall emotional stimuli closely after other emotional stimuli, demonstrating semantic clustering effects around the emotional category (Long et al., 2015; Talmi et al., 2007). The earlier recall of emotional items, as well as the ensuing effect of their early recall on recall of similarly emotional items, neither hinders nor helps them in the emotional pure-list condition. In that condition the test context matches all target stimuli, so it does not help any of the stimuli in particular. But in the mixed list condition the match with context helps emotional stimuli win the competition for recall, while simultaneously hindering the recallability of neutral items. Our model therefore builds on ABC theory and the importance of attention at encoding, but goes beyond it to explain how encoding and retrieval processes interact to result in particular patterns of context-dependent memory performance.

These speculations about the way that context can help us retrieve unique episodic memories tie in well with literature showing that the prefrontal cortex contributes to emotional memory retrieval, and that its contribution can be valence-specific (Dolcos et al., 2012). The prefrontal cortex is of course crucial to retrieval success, with a specific role in the retrieval of emotional memories (Shafer & Dolcos, 2014). Our model can be thought of as depicting the cognitive mechanism that allows this region to support the retrieval of emotional experiences. Of particular relevance for this present manuscript, the prefrontal cortex would have an important role in situations where the retrieval context influences what can be and cannot be recalled. In the mixed list condition the mixed retrieval context hinders recall of a subset of stimuli - the neutral stimuli that were studied together with the emotional ones. Because in that situation memory performance is affected strongly by the text context, encoding activity is a poorer predictor of free recall performance, exactly as Watts et al. (2014) have found when they measured Dm effects. It would be useful to know whether the same findings are obtained when semantic relatedness is controlled.

### Limitations

The experiment reported here has three main limitations. First, because we operationalised emotion using only negative emotionally-arousing stimuli, and therefore we cannot be sure that they generalise for stimuli with positive valence. Although both positive and negative stimuli should capture attention, because this process is thought to be governed by emotional arousal rather than valence (Mather, Clewett, Sakaki, & Harley, 2015; Schimmack & Derryberry, 2005), attention may be allocated differently to stimuli with positive valence. There is evidence, for example, that only negative valence narrows information processing focus (Fredrickson, 2013; Wadlinger & Isaacowitz, 2006), although this difference is less likely to matter to a free recall measure of memory, which captures gist rather than detail. Perhaps most relevant to our work is the finding that compared to negatively-valenced stimuli, positive stimuli led to decreased re-instantiation activity in visual cortex during memory retrieval. Because free recall depends so much on the ability to reinstate studied context (Kark & Kensinger, 2015), if positively valenced stimuli fail in this regard, we may not see enhanced memory for these stimuli in mixed lists.

Second, we have only tested immediate recall here, and therefore cannot be sure that the contextual influences on emotional memory extend in time to delayed tests. It is slightly tricky to replicate the current paradigm in a delayed test, because delayed testing requires carrying out all manipulations between-subjects: participants who study one pure neutral list, one pure emotional list and one mixed list in session 1 cannot be instructed to consider only stimuli from one of these lists in session 2. Thus, unless only a single list is studied in session 1 recall in session 2 would always be mixed. The relationship between immediate and delayed emotional memory effects is of inherent interest to the literature on emotional memory, and warrants additional research.

Finally, unfortunately we could not examine neural indices of successful encoding in our data, because many participants did not have either enough hits or enough misses to analyse the ‘difference due to memory’. It would have been useful to check whether this experiment replicates Watts et al. (2014), and interesting to see whether the effect of emotion on encoding stems particularly from its effect on successful encoding. However, it should be noted that our interpretation of the results does not hinge on those data.

In conclusion, emotional stimuli are prioritised for processing regardless of their local context, but good encoding does not always translate to good memory performance. It is well-known that memory performance depends intimately on the interplay of encoding and retrieval. Our data suggests that this interplay could play an important role in emotional memory enhancement.

## Authors’ note

This study was supported by funding from the Economic and Social Research Council [ES/J500094/1] and was written during DT’s visit at the Princeton Neuroscience Institute. The authors would like to thank S Sunram-Lea and T Sommer for their useful comments; and A Valji, AC Brebenar, and K Mistry for their contributions to stimulus selection, data collection, and data coding. For reprint please contact Deborah Talmi; Division of Neuroscience and Experimental Psychology, University of Manchester, Oxford Road, Manchester, UK.

